# Time-dependent memory of hypoxia exposure influences tumor invasion dynamics

**DOI:** 10.64898/2026.04.07.716866

**Authors:** Gopinath Sadhu, Paras Jain, Ritesh Kumar Meena, Jason Thomas George, Mohit Kumar Jolly

## Abstract

Cancer cells in hypoxic environments often proliferate less but exhibit enhanced migration relative to their normoxic counterparts. Recent *in vitro* and *in silico* studies have characterized the role of hypoxic memory – the ability of cancer cells to retain their hypoxic phenotype even when reoxygenated – in tumor invasion. However, the observations have been limited either to exposing cancer cells to hypoxia for a fixed duration or by assuming a fixed-time persistence of the hypoxic state upon reoxygenation independent of the duration of hypoxia exposure. Thus, time-dependent cell-state changes during hypoxia and their impact on hypoxic memory remains unclear. Here, we first analyze transcriptomic data from breast cancer samples to show that the genes upregulated at transcriptional level and hypomethylated at epigenetic level are enriched in cell invasion, indicating hypoxic memory-driven process of tumor invasion. Next, we used a computational model to investigate how the spatial-temporal dynamics of oxygen levels in a tumor drive time-dependent changes in hypoxic memory and influence tumor invasion dynamics. Our simulation results show that such dynamic hypoxic memory can drive enhanced tumor invasion over a fixed hypoxic memory by a) enriching hypoxic cell density at the tumor front, b) reducing sensitivity of hypoxic cell state to fluctuations in oxygen supply, and c) enhancing effective diffusion of hypoxic cells. Our results highlight the crucial role of dynamic hypoxic memory in shaping tumor invasion dynamics, underscoring the need to elucidate its underlying mechanisms in future studies.

## Introduction

Rapid tumor growth under inadequate vascular supply leads to the development of an oxygen gradient from the blood vessel to distant parts of the tumor, such that interior tumor cells experience a low-oxygen environment (hypoxia) [1, 2]. Cellular responses to hypoxia include up-regulation of hypoxia-associated genes and glycolysis, and down-regulation of cell-cycle genes and oxidative phosphorylation, among others [3, 4, 5]. Thus, depending on the local oxygen concentration, the tumor cells are observed to be in hypoxic or normoxic cell states. Several *in-vitro* studies have highlighted the increased migratory potential of hypoxic cells compared to normoxic cells, and the precise role of hypoxic cells in driving cancer invasion and metastasis is now an active research area *in-vivo* [6, 7, 8, 9]. Recently, the aggressive and metastatic nature of hypoxic tumors was explained by enhancements in the migratory and ROS-resistant characteristics of cells that have experienced hypoxia in the past, i.e., post-hypoxic cells [5, 10]. Post-hypoxic cells were localized around well-oxygenated tumor regions and had higher proportions in circulating tumor cells (CTCs) and distant metastatic lesions than cells that had not experienced hypoxia. Although post-hypoxic cells had an altered molecular phenotype compared with hypoxic cells, they continued to exhibit up-regulated hypoxic hallmark genes, suggesting the presence of hypoxic memory [9, 11]. Such hypoxic memory in post-hypoxic cells was retained even during *in-vivo* metastatic process and *in-vitro* culture under well oxygenated conditions. Based on these observations, Rocha et al.[12] developed an *in silico* tumor model that showed that a sufficiently high duration (50 hours) of persistence time - duration for which post-hypoxic cells maintain their increased migratory behavior after leaving hypoxic regions - was sufficient to explain the presence of post-hypoxic cells in the tumor periphery [12]. Thus, an understanding of the dynamics of hypoxic memory would advance our knowledge of enhanced invasiveness and metastatic ability of hypoxic tumors.

The duration and impact of hypoxic memory can be determined by the extent of phenotypic changes that happen during hypoxic exposure. The initial cellular response to hypoxia is mediated by stabilization of HIF1*α* and activation of its downstream target genes [3]. Moreover, continued hypoxia exposure for multiple days leads to the activation of NF-*κ*B signaling pathways via intracellular H_2_O_2_ ROS generation [5]. Furthermore, other oxygen sensors, e.g., TET DNA demethylases and Jumonji histone demethylases, can act independently of HIF1*α* and cause global changes in DNA methylation and chromatin organization, respectively, contributing to genome-wide expression changes and altered phenotype [13, 14, 15, 16]. Thus, the phenotypic state of cells continues to evolve under hypoxia, with regulation occurring at multiple levels (transcriptional and epigenetic) and across different timescales [17, 18]. It is then imperative to consider that post-hypoxic cells exhibit varying degrees of hypoxic memory depending on their duration of hypoxic exposure. Therefore, a better understanding of hypoxia-driven invasion and metastasis will require us to consider this dynamic nature of hypoxic memory.

In this study, first, by analyzing TCGA breast cancer tumor samples, we show that the exposure to tumor hypoxia can cause persistent cellular invasion through upregulation and hypomethylation of genes driving cell invasion. Then, using our previously developed computational model, we elucidate how spatial-temporal dynamics of oxygen levels within a tumor can cause dynamic changes in hypoxic memory and influence tumor invasion dynamics. The model consists of normoxic and hypoxic cell phenotypes: hypoxic cells can only migrate, whereas normoxic cells proliferate but do not migrate; following the ‘go- or-grow’ dichotomy [19]. Switching between normoxic and hypoxic cell states depends on local oxygen levels, and to incorporate the dynamic nature of hypoxic memory, we considered a time-varying hypoxic-to-normoxic state-switching rate, governed by the duration of hypoxia exposure, with a fixed normoxic-to-hypoxic switching rate. By comparing dynamic hypoxic memory with time-independent hypoxic memory (with a fixed hypoxic-to-normoxic switching rate), we found that dynamic memory accelerated tumor invasion by 1) increasing the proportion of hypoxic cells in the normoxic region and facilitating hypoxic cells at the invasive tumor front and 2) increasing the effective diffusion coefficient of the hypoxic cells. Further, we observed that dynamic memory-driven tumor invasion is less sensitive to the fluctuation in the tumor oxygen supply than fixed hypoxic memory-driven invasion. Overall, our study highlights how dynamic hypoxic memory (or persistence time) can shape the spatiotemporal dynamics of cancer cell invasion.

## Results

### Hypoxia in breast tumor samples epigenetically up-regulate genes contributing to cellular invasion

Recent *in vitro* studies have highlighted how hypoxia promotes cellular migration by inducing epithelial-to-mesenchymal transition via HDAC upregulation [20] and by suppressing DNA demethylases such as TETs [14]. To determine whether epigenetic regulation of cell migration during hypoxia exposure occurs in *in vivo* human tumors, we analyzed primary tumor samples from the TCGA breast cancer patient cohort. The patient cohort of 780 samples, with both transcriptomic and CpG methylation data available, was scored using a chronic hypoxia transcriptional signature identified by analyzing orthotopic tumors in mice bearing MDA-MB-231 cells [10]. The hypoxia scoring of patient samples revealed a unimodal density distribution, from which we selected the bottom 25% and top 25% quartiles and labeled them as normoxic (n = 195) and hypoxic tumor samples (n = 195), respectively (Figure 1a).

**Figure 1:**
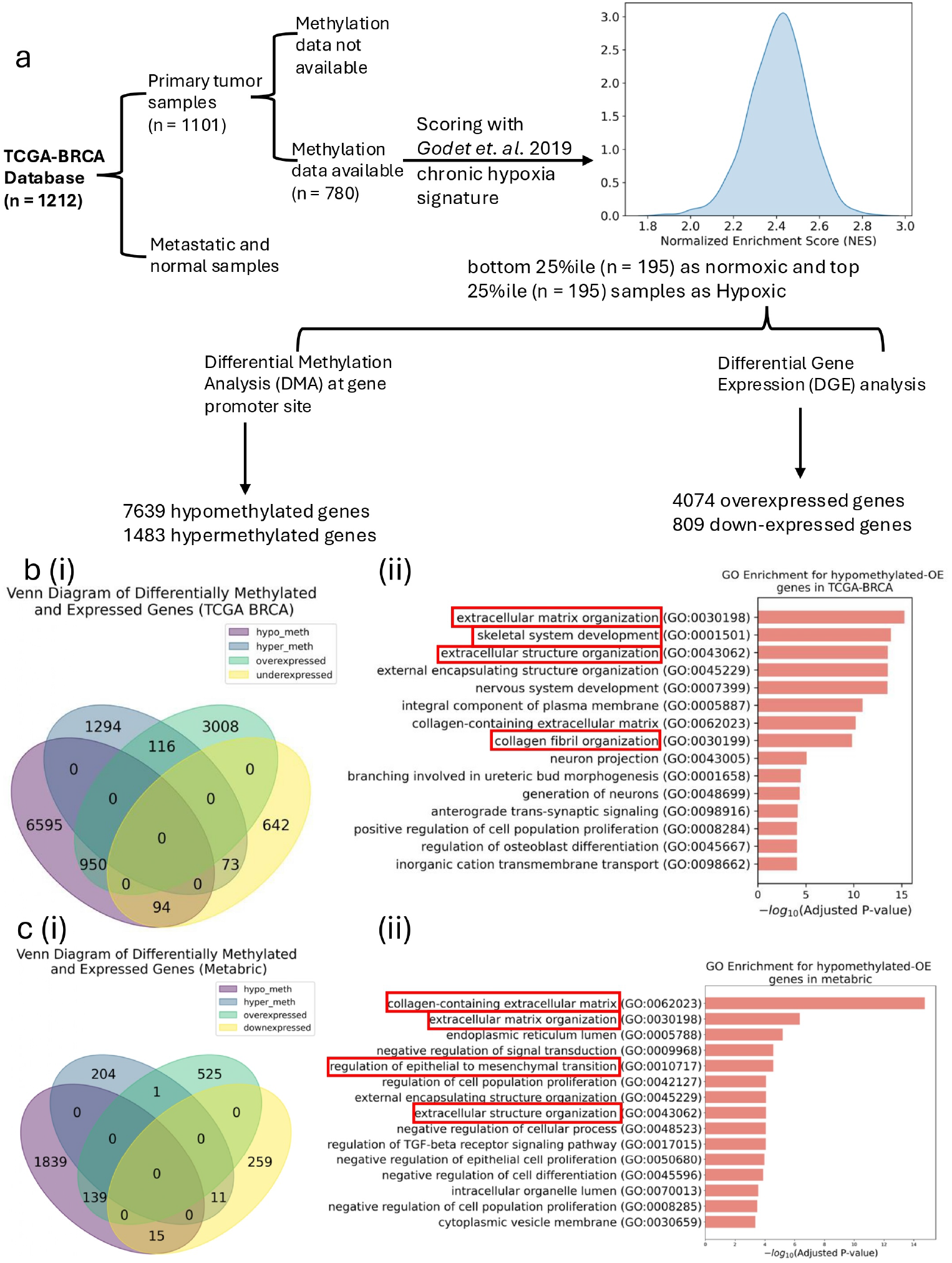
Differential methylation and gene expression analysis of cellular exposure to chronic hypoxia. (a) Pipeline for differential methylation and differential gene expression analysis of TCGA-BRCA dataset to identify gene expression and methylation signatures that significantly differentiate normoxic and hypoxic samples. (b) (i) Venn diagram representing overlap between genes from DMA and DGE analysis on TCGA-BRCA dataset. (ii) Top 15 GO classes (sorted by Adj(p-value)) upon GO analysis of genes that were found overexpressed and hypo-methylated in TCGA-BRCA samples. (c) (i) Venn diagram representing overlap between genes from DMA and DGE analysis on Metabric dataset. (ii) Top 15 GO classes (sorted by Adj(p-value)) obtained upon GO analysis of genes that were found overexpressed and hypo-methylated in METABRIC samples.

Next, using the above-labeled normoxic and hypoxic tumor samples, we identified differentially regulated genes both at the transcriptional and epigenetic levels in response to chronic hypoxia. While differential gene expression analysis revealed that 4074 genes were over-expressed and 809 genes were down-regulated, differential methylation analysis revealed that 7639 genes were hypo-methylated and 1483 genes were hyper-methylated (Figure 1a). Next, to check for the overlap between differentially methylated and differentially expressed genes, we constructed a venn diagram with the four classes of genes: 1) hypermethylated and over-expressed, 2) hyper-methylated and down-regulated, 3) hypo-methylated and over-expressed, 2) hypo-methylated and down-regulated. We found that the number of genes that were both hypomethylated and overexpressed was much higher (n = 950) than other combinations (Figure 1b,i). Thus, we next studied the functional relevance of both hypo-methylated and over-expressed genes by performing gene ontology (GO) analysis. Intriguingly, the analysis revealed a strong enrichment of GO classes related to ECM organization, extracellular structure organization, and other cellular invasion-related classes (highlighted in red boxes in Figure 1b,ii and Supplementary Table 1 & 2).

To further validate our results above, we performed similar analyses using another publicly available breast cancer patient dataset, the METABRIC dataset [21]. Importantly, we observed the same trends again: a) the hypo-methylated and over-expressed genes were in greater numbers, and b) GO classes related to cellular invasion were enriched in the METABRIC dataset as well (Figure 1c and Supplementary Table 4). Thus, our analyses above highlight the epigenetic regulation of cellular invasion in patient tumor samples.

Next, to test whether differentially hypo-methylated and over-expressed cellular invasion-related genes from breast cancer hypoxic tumor samples show significant up-regulation in independent analyses of hypoxic exposures as well, we constructed a gene signature ‘*HOI*’ (**Hypo**-methylated; **Over**-expressed and **Invasive**) using the genes that were hypo-methylated, over-expressed, and involved in cell invasion GO terms. This gene set was used to score the publicly available cell line transcriptomic datasets via ssGSEA [22]. The transcriptomic analysis of MDA-MB-231 (GSE292924) [23] (a triple-negative breast cancer cell-line) exposed to hypoxia for 48 hours did not show statistically significantly increased activity level of the *HOI* genes (p-val=0.07, Figure S1 ai), indicating that 48 hrs of hypoxia exposure was perhaps insufficient for epigenetic alteration of cell invasion genes. Thus, we next analyzed two publicly available transcriptomic datasets where cells were exposed to hypoxia for longer (up to 72 hr): 1) ovarian cancer cell line, SK-OV-3 (GSE53012)[24], and 2) Human umbilical cord mesenchymal stem cells, hUC-MSCs (GSE298520) [25]. Both cell lines showed a strong increase in the activity levels of the *HOI* gene set (p-value *<* 0.01) (Figure S1 aii, aiii). Further, GO classes of differentially expressed genes between 0 and 72 hr time points in the above two cell lines were similar to what we observed in our analysis of normoxic vs hypoxic tumor samples (Figure S1 bi, bii, Figure 1bii and Supplementary Table 5 and 6). These independent observations in cell lines further support our hypothesis that, when cancer cells experience prolonged hypoxic conditions, epigenetic changes occur which regulate the expression of a set of genes that promote cellular invasion and ECM degradation. Further, since the *HOI* signature derived from breast cancer tumor samples also shows consistent trends in other cancer types too, these results support the notion that hypoxia-induced cellular invasion via epigenetic regulation may be a more universal mechanism by which cancer cells regulate their gene expression in response to chronic hypoxia.

### Dynamic hypoxic memory accelerates tumor invasion by enriching hypoxic cells at the tumor front

We observed significant epigenetic regulation of genes that contribute to cellular invasion during hypoxic exposure. Given that these epigenetic changes can persist for longer, even upon re-exposure to normoxic conditions, post-hypoxic cells continue to invade their surroundings. However, because induction and erasure of epigenetic changes are time-dependent processes, we next used a computational model to study how dynamic changes in hypoxia memory influence tumor invasion dynamics.

We extended our previous computational model to study tumor invasion with dynamic hypoxic memory in spatially heterogeneous oxygenated conditions [19]. Specifically, we have considered that, at time *t* = 0, a population consisting of normoxic cells at the center of the spatial domain (4≤ *x*≤ 6), and normoxic cells surrounded by extracellular matrix (ECM) (0≤ *x <* 4 and 6 *< x*≤ 10) (Figure 2a). Oxygen is supplied from the domain boundaries (at *x* = 0 and *x* = 10) and is uniform across the spatial domain initially (*t* = 0). As the population grows with time, an oxygen gradient is established and oxygen concentration around the tumor center drops below the threshold levels (*c*_*H*_), triggering normoxic-to-hypoxic state transitions. The resulting hypoxic cells migrate in their neighborhood by diffusion and haptotaxis, degrade the surrounding ECM and facilitate further tumor growth by reverse transitioning into the normoxic state under well oxygenated conditions (*c > c*_*H*_). Now, to study the role of hypoxic memory in tumor invasion, we considered that when hypoxic cells get re-oxygenated by migrating into the normoxic region, they transition to the normoxic state at a rate defined by the parameter *µ*_*hn*_. For fixed hypoxic memory durations (i.e. fixed persistence time [12]) *µ*_*hn*_ is set to a constant value, *µ*_*hn*_ = *µ*_*hn*,0_. For dynamic hypoxic memory, we consider time-dependent changes in *µ*_*hn*_ based on the local oxygen levels (*c*), basal transition rates (*µ*_*hn*,0_), and the memory induction/erasing timescales (*β*) (Eq. (3)). Such time-dependent variation in *µ*_*hn*_ leads to slower hypoxic-to-normoxic switching for an increasing duration of past hypoxic exposure (Figure 2b).

**Figure 2:**
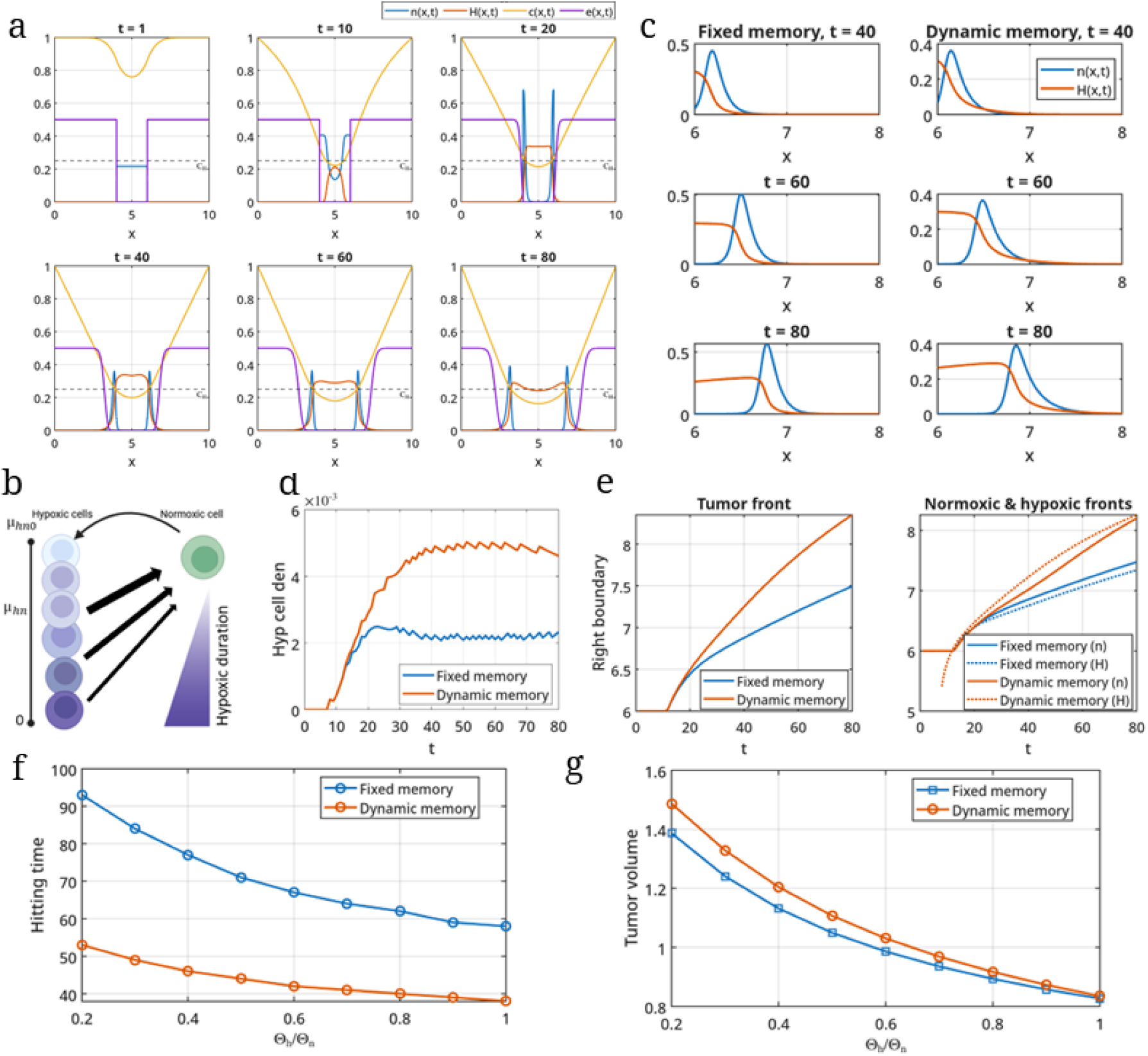
Dependence of hypoxic memory on the duration of hypoxia exposure enhances tumor invasion. (a) The spatial distributions of normoxic cell (*n*(*x, t*)), hypoxic cell (*h*(*x, t*)), ECM (*e*(*x, t*)) densities and oxygen concentration (*c*(*x, t*)) at *t* = 1, 10, 20, 30, 40, 50, 60, 70 and 80 with dynamic hypoxic memory. (b) Schematic depicting the phenotypic transitions between normoxic and hypoxic phenotypic states with constant rate of normoxic-to-hypoxic transition, and the extent of hypoxia exposure dependent rate of hypoxic-to-normoxic transition. The thickness of the arrow highlight relative rates of hypoxix-to-normoxic state transition. (c) The spatial distribution of normoxic and hypoxic cell densities at the tumor periphery at *t* = 40, 60, and 80. (d) The hypoxic cell density in the normoxic region, where oxygen concentration *> c*_*H*_ changes over time. (e) Comparison of the time evolution of the tumor right boundary, and normoxic and hypoxic front propagation. Here, the the right boundary, hypoxic and normoxic boundaries being the spatial locations *x*, where *n*(*x, t*) + *h*(*x, t*) = 10^*−4*^, *h*(*x, t*) = 10^*−*4^ and *n*(*x, t*) = 10^*−*4^, respectively for given *t*. The effects of fold change of oxygen consumption rate of hypoxic cell with respect to the consumption rate of normoxic cells on (f) tumor invasion and (g) tumor volume. In all the analyses above, the period of oscillation is 30 and 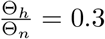 unless stated otherwise.

Numerical simulations of tumor growth and invasion showed that dynamic hypoxic memory significantly increased the proportion of hypoxic cells in the normoxic region when compared to the fixed memory scenario (Figure 2c, d). Furthermore, hypoxic cells led the tumor front in case of dynamic memory but not with the fixed memory (Figure 2e). Therefore, we observed that hypoxia-driven tumor invasion was faster with dynamic memory than fixed memory (Figure 2e). The enhanced invasion with dynamic memory relative to fixed memory remained consistent even when slower basal switching rates (*µ*_*hn*,0_) were considered for both memory modes (Figures S1a and S1b). However, with smaller *µ*_*hn*,0_, even fixed hypoxic memory led hypoxic cells to lead the tumor front, but only during the initial phases of growth.

Hypoxic cells and normoxic cells often have different metabolic dependencies - the former usually rely on glycolysis, whereas the latter depend on oxidative phosphorylation; thus, hypoxic cells often consume oxygen at a slower rate than normoxic cells [4]. Such differences in oxygen consumption between the two phenotypes shape the oxygen gradient within the tumor, modulate the extent of the hypoxic region, and thus may impact tumor invasion. Therefore, we next investigated how the differences in oxygen consumption rates between the normoxic and hypoxic phenotypes affected tumor invasion and growth dynamics with dynamic and fixed hypoxic memory. On varying the consumption rate of hypoxic cells relative to normoxic cells, 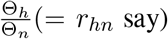, we found that for fixed hypoxic memory, there was a significant decrease in tumor volume and slower invasion as the Θ_*h*_ ranges from being 20% of Θ_*n*_ to being equal to Θ_*n*_ (Figure 2f, g).

Tumor volume reduces because the oxygen gradient in the tumor core establishes rapidly for larger *r*_*hn*_ values (Figure S1c). A similar drastic change in tumor volume with increasing *r*_*hn*_ values was observed with dynamic hypoxic memory as well; however, tumor invasion was only slightly affected (Figure 2f, g). These results highlight how large differences in oxygen consumption rates between normoxic and hypoxic cells, along with dynamic hypoxic memory, lead to tumors that grow larger and invade more rapidly.

### Dynamic hypoxia memory-driven invasion is less sensitive to fluctuations in tumor oxygen supply

Tumor oxygenation from the nearby blood vessels could vary over different timescales: a) short timescales (hours) due to fluctuations in flux of red blood cells from inefficient vasculature, and b) long timescales (days) due to angiogenesis and vascular remodeling, with two-to-five fold differences between the low and high oxygenated levels [26, 27]. Thus, we next examined how time-varying oxygenation leads to time-varying changes in tumor hypoxia, and thereby affects tumor volume and invasion with fixed and dynamic hypoxic memory. To implement this setup, we periodically fluctuated the oxygen supply at the domain boundaries (*x* = 0 and *x* = 10) between the two levels *c*_high_ and *c*_low_, with a period *T*. Figure 3a shows temporal changes in the oxygen gradient and the hypoxia region (*c < c*_*h*_) along with cell densities, as the oxygen supply varies between *c*_high_ = 1 and *c*_low_ = 0.8 with a period of 30 units. The above temporal changes in oxygen gradient, hypoxia region and resulting cell densities, were significantly different from tumor growth under constant oxygenation (*c*_high_ = *c*_*low*_ = 1) (Figure 3b); with fluctuating oxygenation leading to faster tumor invasion but with smaller tumor volume for both the memory modes (Figure 3a, b). Next, we specifically analyzed the effects of fluctuation amplitude and period on tumor invasion and growth across the two memory modes.

**Figure 3:**
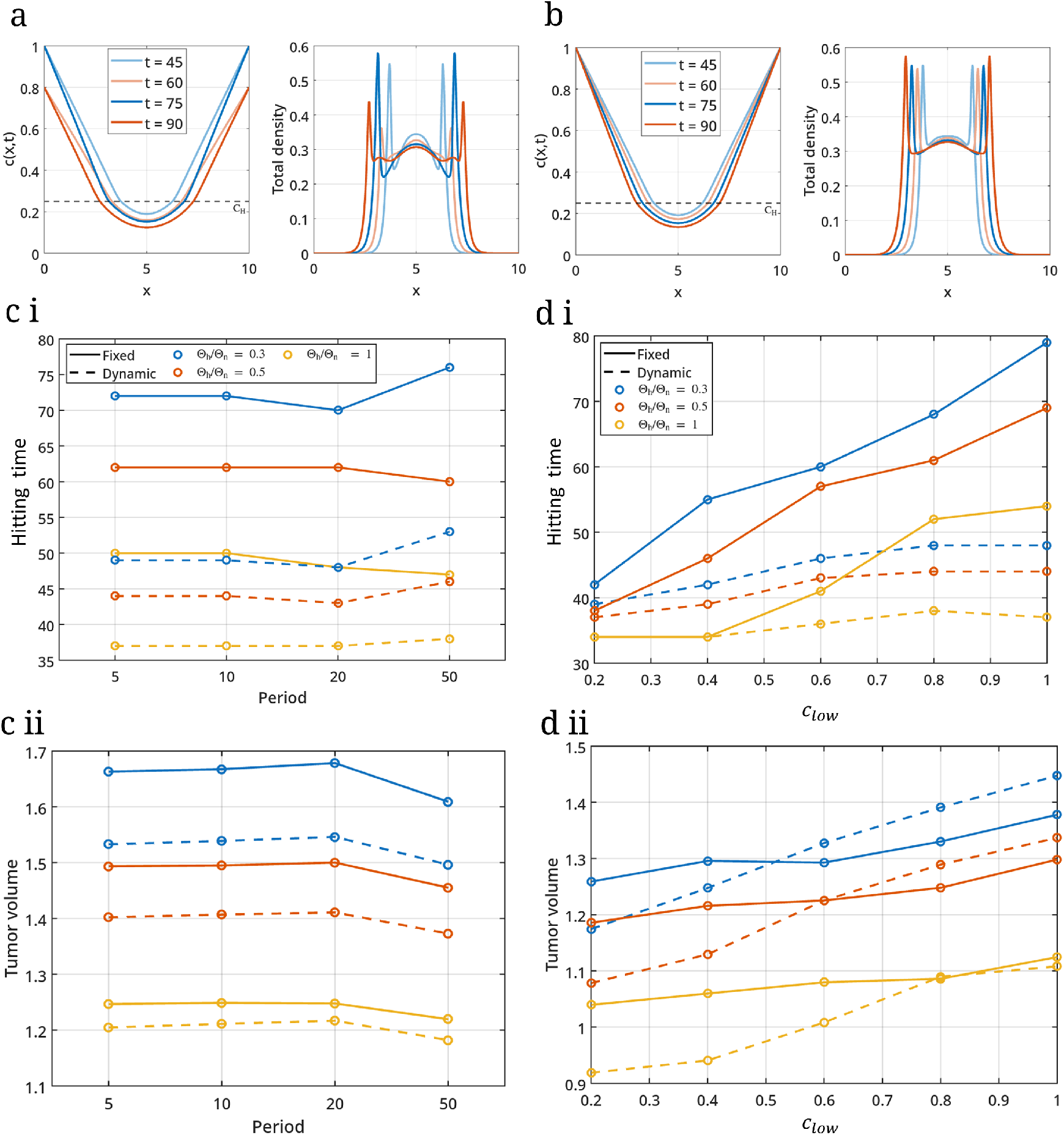
Fluctuations in tumor oxygenation from the boundary further enhances invasion, for both fixed and dynamic hypoxic memory scenarios. Spatial-temporal dynamics of oxygen distribution and total tumor cell density (sum of normoxic and hypoxic cell densities) when the tumor oxygenation from the boundary is (a) oscillating with period 30 time units, and (b) constant. The sampled time points, *t* ∈ {45, 75} highlight oxygen profiles and cell densities by the end of high oxygenation condition (*c*_high_) of duration 15 units, and *t*∈ { 60, 90} highlights oxygen profiles and cell densities by the end of low oxygenation condition (*c*_low_) of duration 15 time units. (c) Time to reach a spatial location 7.5 and tumor volume at *t* = 100 for different periods of oscillation in tumor oxygenation at the tumor boundary for varying ratio of oxygen consumption rates, *r*_*hn*_. (d) The effects of amplitude of oscillation on time to reach *x* = 7.5 and tumor volume at *t* = 100 for varying *r*_*hn*_ levels. In all the analyses above, the period of oscillation is 30, *c*_high_ = 1 and *c*_low_ = 0.8, and *r*_*hn*_ = 0.3 unless stated otherwise.

By fixing the amplitude of fluctuation and varying the period lengths from 5 units (half of the cell doubling time) to 50 units (five times the cell doubling time), we observed only slight changes in tumor volume and invasion speed (Figure 3ci, cii). With dynamic memory, we observed that tumor volume and invasion speed peak at a period length of 20 units, irrespective of relative oxygen consumption rates (*r*_*hn*_). However, the fixed-memory optimal period was observed to strongly depend on the *r*_*hn*_ value (corresponding to changes in the curvature of the solid lines in Figure 3ci for increasing period length, from non-monotonic for *r*_*hn*_ = 0.3, to non-increasing for *r*_*hn*_ = 0.5 and monotonically decreasing for *r*_*hn*_ = 1).

Upon fixing the period of fluctuation, we observed that fluctuations of larger amplitudes (smaller *c*_low_ values) increased tumor invasion, irrespective of the hypoxic memory type (fixed or dynamic) and the oxygen consumption rates of normoxic and hypoxic cells (Figure 3di). Moreover, invasion driven by fixed hypoxic memory was more sensitive to amplitude variation than invasion driven by dynamic hypoxic memory. However, this trend in the sensitivity of the two memory modes to oxygenation amplitude variation was reversed when tumor volume changes were observed (Figure 3dii). We found significant decreases in tumor volume with decreasing *c*_low_ values in the dynamic memory case compared to the fixed memory case. Further, the optimality of dynamic memory over fixed memory, resulting in larger tumor volume and faster invasion, remained intact up to a threshold fluctuation amplitude (*c*_low_ values corresponding to the crossover of the dashed and solid lines of the same color in panel Figure 3dii). The threshold amplitude depended on the oxygen consumption rate ratio between hypoxic and normoxic cells, *r*_*hn*_.

Overall, we observed that dynamic hypoxic memory-driven invasion was less sensitive to periodic changes in tumor oxygenation than fixed hypoxic memory-driven invasion. Further, we consistently observed that dynamic memory led to faster invasion than fixed memory and, in some cases, to larger tumor volume as well.

### Dynamic hypoxia memory enhances the effective diffusion of hypoxic cells

In the previous section, we observed that dynamic memory enhanced tumor invasion by enriching hypoxic cells at the tumor front. To investigate further how the two memory modes drive invasion, we compared their influence on 1) haptotaxis and 2) cell diffusion, the two modes of hypoxic cell migration in our model setup. For the comparison, we quantified the dynamic changes in coefficients of haptotaxis (ECM gradient, 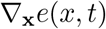 and effective diffusion (*D*_*h*_(1 *−n*(*x, t*) *−H*(*x, t*) *−e*(*x, t*))) at the tumor periphery, defined by 0.2-unit distance from the tumor front.

Upon first investigating tumor invasion under constant oxygen supply from the boundary, we found that although the two memory modes had similar haptotaxis coefficients over simulation time, the effective diffusion coefficient was higher in the dynamic memory condition (Figure 4a, b). Further, dynamic memory had a higher effective diffusion coefficient than fixed memory despite having a larger tumor volume, an outcome of a dispersed (spread of cell density) migration mode (Figure 2b, g). Moreover, as dynamic memory led to more hypoxic cells in both the normoxic region and the tumor front, the overall resulting haptotaxis and cell diffusion were significantly higher with dynamic than with fixed memory (Figure 3di, Figure 4c).

**Figure 4:**
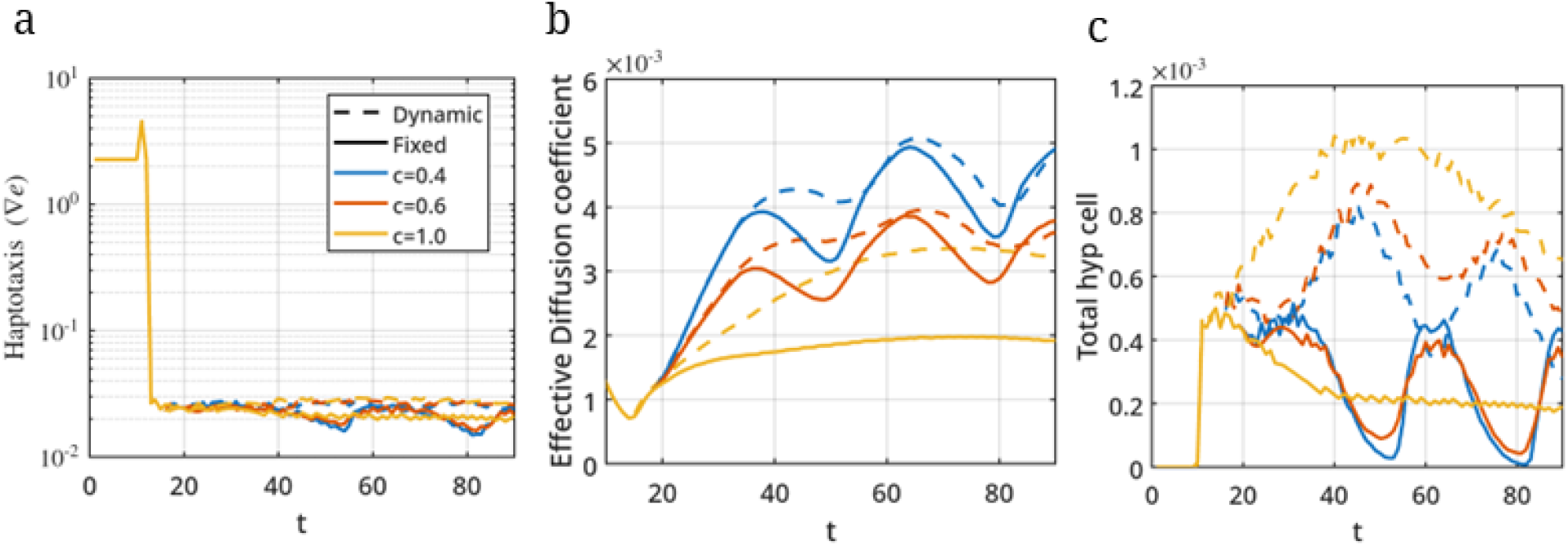
Higher effective diffusion of hypoxic cells with dynamic hypoxic memory. Time evolution of a) haptotaxis coefficient (ECM gradient), and (b) effective diffusion coefficient, at the tumor periphery for different values of *c*_low_. (c) Dynamics of tumor hypoxic cell densities at the tumor periphery for different oscillation periods of tumor oxygenation. Here, the tumor periphery is the region lying 0.2 spatial distance inside the tumor front. In all the analyses above, the period of oscillation is 30, *c*_high_ = 1, and 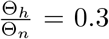, unless stated otherwise.

With fluctuating tumor oxygen supplies, we observed that, while the haptotaxis coefficient (ECM gradient) at the tumor periphery remained unchanged across both memory modes with increasing fluctuation amplitude, there was a significant average increase in the effective diffusion coefficient (Figure 4a, b, Figure S4b, c). Such an increase in the effective diffusion coefficient results from reduced tumor volume, especially reduced normoxic cell density, because of insufficient oxygen for their sustenance and growth (Figure 3dii). Further, while the effective diffusion coefficient was always higher for dynamic memory, it was less sensitive to oxygen fluctuations as well, because of reduced transitions from hypoxic to normoxic states (Figure 4b). Oxygen fluctuations and the concomitant reduction in tumor volume, on the other hand, also reduced overall hypoxic cell density at the tumor periphery in both memory modes; however, hypoxic cell density remained higher with dynamic memory (Figure 4c). Thus, dynamic memory maintaining 1) a similar haptotaxis coefficient, 2) a higher and persistent diffusion coefficient, and 3) higher hypoxia at the tumor periphery as compared to the fixed memory mode collectively drove faster invasion.

## Discussion

Tumor hypoxia is linked to aggressive disease and poor prognosis. The enhanced metastatic ability of hypoxic tumors has been linked to persistent invasiveness, ROS-resistance, and an immune evasion state of the tumor cells that were exposed to prolonged hypoxia [10, 5, 11]. The above sustained phenotypic changes in post-hypoxic tumor cells have been previously referred to as ‘hypoxic memory’ and can occur because of cellular adaptation to hypoxia exposure at multiple levels of gene regulation – transcription, chromatin reorganization, and DNA methylation – over different timescales, from minutes to days. Although a few *in vitro* and *in silico* studies have characterized the role of post-hypoxic cells in tumor invasion, the observations have been limited to cases where either 1) cancer cells have been exposed to hypoxia for a fixed duration or 2) the persistence time in the hypoxic (invasive) cell state upon re-oxygenation is independent of the duration of hypoxia exposure [9, 12]. Thus, the duration of hypoxia exposure and its influence on the persistence of invasive traits among post-hypoxic cells, and their overall impact on tumor invasion, are largely unknown. Here, by first analyzing TCGA breast tumor samples, we showed that tumor hypoxia exposure can enable enhanced cellular invasion through coordinated transcriptional and epigenetic changes for relevant genes, and then we investigated how the dynamics of the above hypoxic-induced epigenetic changes can affect tumor invasion by using our previously developed framework [19].

Our differential methylation analyses between normoxic and hypoxic breast tumor samples showed that nearly 7700 genes were hypomethylated, about 5 times as many as were hypermethylated under hypoxic exposure (Figure 1bi, ci). Such extensive global hypomethylation under hypoxia has been reported in earlier studies as well [28, 13, 29]. Further, the genes that were hypo-methylated and over-expressed contributed to cellular invasion, as they were part of EMT and ECM reorganization cellular programs, as compared to hyper-methylated and down-expressed genes whose association with cellular invasion was non-significant (Figure 1bii, cii, Supplementary Table 1 and 3). The presence of epigenetic regulation of pro-invasive genes but not of anti-invasive genes suggests that activation of pro-invasive genes can persist longer, even upon reoxygenation, when anti-invasive genes are also reactivated. Such simultaneous expression of both pro- and anti-invasive genes can lead to distinct modes of tumor invasion [30, 31, 32, 33]. We note that our analysis of epigenetic regulation is limited to changes in DNA methylation; however, epigenetic regulation also occurs through chromatin remodeling via histone modification, and anti-invasive genes may be regulated more by histone modifications during hypoxia [15, 16]. Joint analysis of DNA methylation and histone modification data can help us better understand hypoxia-induced epigenetic regulation of cellular invasion.

We currently lack quantitative experimental data on how the hypoxic-to-normoxic phenotype switching rate, *µ*_*hn*_ alters during hypoxia and reoxygenation. Thus, to model a dynamic hypoxic memory, we chose the timescales of *µ*_*hn*_ changes based on the existing knowledge of memory timescales for other well-characterized cellular processes [17, 18, 34, 35, 36]. Thus, during normoxia exposure *µ*_*hn*_ increases on timescales at which chromatin-based epigenetic markers get erased, which is about the duration of one to two cell divisions (see *β* parameter in Eq. 3 and Table 1). Further, since existing experimental reports captured chromatin- and DNA-based epigenetic changes after short-term hypoxia exposure, we also assumed *µ*_*hn*_ to decrease on timescales of one to two cell divisions during hypoxia exposure [9, 14, 15, 16] (see *β* parameter in Eq. 3 and Table 1).

**Table 1:** Table of the model parameters appearing in equations (1)-(5)

| Parameter | Description | Dimensional value | Dimensionless value | Source |
| --- | --- | --- | --- | --- |
| $D_h$ | Diffusivity of hypoxic cells | $5 \times 10^{-10} - 2.5 \times 10^{-9} \text{ cm}^2 \text{ s}^{-1}$ | 0.01 | [54] |
| $\lambda_n$ | Proliferation rate of normoxic cells | $0.014 - 0.056 \text{ hr}^{-1}$ | 0.1 | [55] |
| $\mu_{nh}$ | Transition rate from normoxic to hypoxic | $1 \text{ hr}^{-1}$ | 0.5 | [56] |
| $\gamma_n$ | Necrotic death rate of normoxic cells | $\frac{\lambda_n}{10}$ | 0.001 | [19] |
| $\gamma_h$ | Necrotic death rate of hypoxic cells | $\gamma_n$ | 0.001 | [19] |
| $\delta$ | ECM degradation rate | — | 15 | [19] |
| $c_H$ | Hypoxic threshold value of oxygen | $2.5 \text{ mmHg}$ | 0.25 | [55] |
| $c_N$ | Necrotic threshold value of oxygen | $1 \text{ mmHg}$ | 0.1 | [55] |
| $\xi_h$ | Haptotaxis coefficient | — | 0.001 | [19, 57] |
| $\alpha$ | Strength of memory induction | — | 0.8 | [19] |
| $\beta$ | Timescale of memory induction/erasing | — | 10 | [19] |
| $\mu_{hn0}$ | Basal transition rate of hypoxic to normoxic | $\frac{1}{96} \text{ hr}^{-1}$ | 0.5 | [56] |

Under constant tumor oxygenation at the spatial domain boundary, we found that tumor invasion driven by dynamic hypoxic memory was faster than that driven by fixed hypoxic memory (Figure 2e). Such enhanced invasion with dynamic hypoxic memory was due to a greater presence of hypoxic cells in the normoxic region compared to fixed memory, with hypoxic cells leading the tumor front in the dynamic but not fixed memory scenario (Figure 2d, e). Similar presence of hypoxic (or post-hypoxic) cells at the tumor periphery was also observed in breast tumors originating from orthotopic mouse implantation of MDA-MB-231 cells and transgenic mouse models, and was associated with enhanced tumor invasion [10]. Further, in the above experiments, when the post-hypoxic cells (hypoxic cells in the oxygenated regions) from the tumors were sorted and grown under ideal (21%) oxygen environment *in vitro*, they retained their original phenotypic state. However, in a control experiment, MDA-MB-231 cells lost the hypoxic state they had attained during two days of hypoxia exposure when returned to normoxic conditions [10]. Thus, the above observations suggest that post-hypoxic memory effects are stronger upon prolonged hypoxia exposure in tumors, which reduces their reversal rate to the normoxic state upon re-oxygenation. Our model captured this stabilization of hypoxic cells in the well-oxygenated regions of the tumor as we observed high hypoxic cell densities in the normoxic regions, concentrated around small *µ*_*hn*_ values (Figure S5).

Under periodic changes in tumor oxygenation at the spatial domain boundary, we observed enhanced tumor invasion with increasing oscillation amplitude and intermediate oscillation period (timescales of two cell divisions; Figure 3c, d) with both fixed and dynamic memory modes. Numerous *in vitro* studies across cancer types have reported that cells exhibit enhanced invasiveness and migratory traits when exposed to fluctuating hypoxia compared to chronic hypoxia [37, 38, 39]. Further, tumor-bearing mice, when exposed to fluctuating oxygen levels in their environment, indicate a positive association between fluctuating hypoxia and metastatic potential of tumors [40, 41, 42, 43]. For example, when tumor-bearing mice, with tumors originating from orthotopic implantation of human cervical cancer cells, were exposed to fluctuating oxygen levels between 5% and 20% periodically every 10 minutes for 12 cycles/day and for 21 days, they had greater lymph node metastasis than control mice exposed to ambient (21%) oxygen conditions throughout the experiment [41]. Moreover, these cyclically oxygenated mice had smaller tumors, a possible outcome of reduced tumor growth due to limited oxygen availability (Figure 3dii). Furthermore, when human tumor samples were stratified using a cycling hypoxia gene signature derived from cycling MDA-MB-231 cells under hypoxic and normoxic conditions, samples positive for the cycling hypoxia signature had poor disease prognosis [44]. We note that the above associations are observed between fluctuating hypoxia and metastases, in which tumor invasion is just an initial step; more careful histopathological and molecular analyses of primary tumor samples that have experienced fluctuating hypoxia can help us better characterize its effects on tumor invasion and decipher the dynamic vs fixed nature of hypoxic memory.

While quantifying the contributions of haptotaxis and cell diffusion in enhancing tumor invasion with dynamic memory relative to fixed memory, we observed that dynamic memory results in a higher effective diffusion coefficient for hypoxic cells at the tumor periphery than fixed memory, while there was no significant difference in the haptotaxis coefficient (ECM gradient). Moreover, although fluctuations in tumor oxygen supply reduced hypoxic cell density at the tumor periphery, the significant increase in the effective diffusion of hypoxic cells led to faster invasion with increasing amplitude of oxygen fluctuations. Thus, tumor invasion is a complex interplay between hypoxia memory and the spatial-temporal profile of oxygen, regulated by continuous tumor growth and fluctuations in oxygen supply. Although we have considered the independence of oxygen fluctuations and the formation of an oxygen gradient from tumor growth, they are causally related. Formation of a hypoxic core triggers angiogenesis and leads to the formation of new, but inefficient, vasculature with highly irregular oxygen supply [45], and such cross-talk could enhance tumor invasiveness over time [46, 47].

Overall, in this study, we have investigated how spatio-temporal changes in oxygen levels within a tumor encode a dynamic memory of past hypoxic exposures, influencing the tumor invasion dynamics. However, tumor invasion depends on other microenvironment factors as well, e.g., the mechanical properties of the extracellular matrix. Cells grown under cyclic changes in ECM stiffness encode mechanical memory of past ECM stiffness, contributing to their distinct invasion characteristics [48, 49]. Combining these distinct phenomena in a single modeling framework would provide a detailed description of tumor invasion dynamics. Further, recent *in vitro* and *in vivo* observations highlight the interconnection between the hypoxic cell state and immunoresistance via downregulation of IFN signaling, and between the hypoxic cell state and chemo- and radio-resistance via enhanced HIF1*α* signaling and ROS resistance. Modeling such hypoxia-induced switching to resistant states can help us understand how tumor microenvironment factors modulate treatment outcomes and may help us design better therapeutic strategies [50, 51].

## Methods

### Segregating tumors into normoxic and chronic hypoxia tumors

Data for TCGA-BRCA samples was obtained from the UCSC-Xena browser and filtered for three criteria: (1) The sample must be a primary tumor sample; (2) The sample must have gene expression data available; (3) The sample must have methylation data available. These filters resulted in 780 samples which were then scored for chronic hypoxia using a 41-gene signature previously identified [10]. The samples were then ranked based on their ssGSEA score for the signature and segregated into three classes: samples with expression profile similiar to (1)First quartile: Normoxic tumors (n = 195), (2) Fourth quartile: Hypoxic tumors (n = 195) (3) Intermediate hypoxic, (n = 390).

Similar analysis done on METABRIC data (obtained from cbioportal) samples (n = 1420) identified: (1) 355 Normoxic tumors; (2) 355 Hypoxia tumors, and (3) 710 Intermediate hypoxia tumors.

### Differential methylation analysis

The aim of this analysis was to figure out genes whose promoter methylation status is significantly altered in the normoxic and chronic hypoxia tumor samples. The analysis was performed using an R package, limma [52] for both TCGA-BRCA and METABRIC datasets.

In the TCGA-BRCA samples for statistical filtering, CpG sites with Adj(P.Value) *<* 0.05 and (logOdds ratio) *>* 0.1 were filtered and considered for downstream analyses. CpG sites were then mapped to respective genes and the CpG sites within 2500 bp upstream to 500 bp downstream of the transcription start site (TSS) were considered to be lying over the promoter region of the gene. Next, to determine the overall hypo-/hyper-methylation status of gene promoters, we first filtered out promoters with fewer than three significantly altered CpG sites. Promoters with a ratio

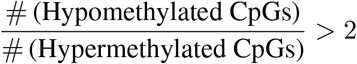

were labeled as hypomethylated, whereas promoters with

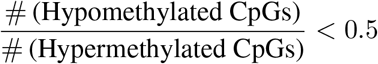

were labeled as hypermethylated.

Similarly, in METABRIC samples again, CpG sites with Adj(P.Value) *<* 0.05 and (logOdds ratio) *>* 0.1 were filtered. Filtering based on the number of CpGs per promoter site was performed only for TCGA samples, as METABRIC provided methylation data for only one CpG site per gene promoter.

The analysis revealed:

- 7639 genes were hypomethylated genes in TCGA-BRCA.
- 1483 genes were hypermethylated genes in TCGA-BRCA.
- 1993 genes were hypomethylated in METABRIC.
- 216 genes were hypermethylated in METABRIC.

#### Differential Gene Expression analysis

The aim of this analysis was to identify genes that were differentially expressed in normoxic and chronic hypoxia tumor samples. The differential expression was performed using limma [52]. Upon DGE analysis genes were filtered based on (1) Log(FC) *>* 1 and Adj(P.Value) *<* 0.05in TCGA-BRCA and (2) Log(FC) *>* 0.5 and Adj(P.Value) *<*0.05 in METABRIC. The analysis revealed:

- 4074 genes were overexpressed in TCGA-BRCA.
- 809 genes were downexpressed in TCGA-BRCA.
- 665 genes were overexpressed in METABRIC.
- 285 genes were downexpressed in METABRIC.

### ssGSEA and GO analysis

Publicly available gene expression datasets were obtained from NCBI GEO website, namely, GSE292924, GSE53012, GSE298520. Different preprocessing steps were carried for RNA-seq and Microarray datasets. For RNA-seq datasets, the respective count matrices were obtained from NCBI GEO and converted to TPM, which were log-transformed and used for downstream analyses. For microarray datasets the respective raw “.CEL.gz” files were downloaded from NCBI GEO and Robust Multiarray Average (RMA) normalisation was performed to obtain normalised intensity data matrix with rows as genes and columns as samples. Single-sample GSEA (ssGSEA), an extension of Gene Set Enrichment Analysis (GSEA) [22], calculates separate enrichment scores for each pairing of a sample and a gene set. Each ssGSEA enrichment score represents the degree to which the genes in a particular gene set are coordinately up- or down-regulated within a sample.

For bulk RNA-seq and microarray datasets, `GSEApy` [53], a Python package, was used to perform both ssGSEA and GO analysis.

#### Making the HOI gene set

The aim of this analysis was to identify genes that were: (1) Significantly hypomethylated at the promoter region; (2) Significantly overexpressed at the RNA level; (3) Directly or indirectly promote cellular invasion, ECM degradation, or EMT. Only the TCGA-BRCA dataset was utilised for this analysis, as information about methylation status of more CpG sites (approx 450K) was available as compared to just one CpG site per gene (approx 13k) in METABRIC. After filtering for methylation and expression status, we conducted GO analysis on the resulting gene set. GO classes were further filtered based on Adj(p.value) *<* 0.001 (Supplemetary table 3). Lastly, GO classes that were related to promoting cellular invasion, ECM degradation, or EMT were picked, and a union of genes in these classes was taken to construct the invasion gene set. (The geneset identified is provided in Supplementary table 7).

### Mathematical model

We consider a mathematical model of tumor growth that adheres to the go-or-grow dichotomy. Here, normoxic cells follow grow property as they are highly proliferative. On the other hand, hypoxic cells exhibit a go property, as they are highly migratory. The rates of phenotype switching between these cell types depend on the local oxygen levels within the tumor microenvironment. Let *n*(**x**, *t*) denotes the density of normoxic cells and *h*(**x**, *t, µ*_*hn*_) represents the density of hypoxic cells. Additionally, *c*(**x**, *t*) denotes the concentration of oxygen. Here, **x** = (*x, y, z*) indicates spatial coordinates, *t* represents time, and *µ*_*hn*_ is a variable in the phenotypic space 𝒫= [0, *µ*_*hn*0_], where *µ*_*hn*0_ indicates the basal transition rate from the hypoxic to the normoxic phenotype. The density of hypoxic cells across the phenotypic space is given as

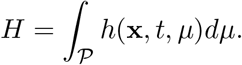

The normoxic cell density dynamics is given as [19]

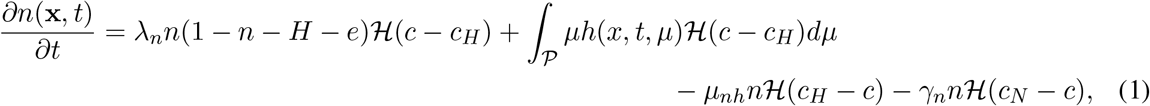

where *ℋ* denotes the Heaviside function and is defined as

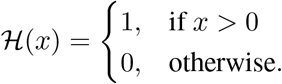

The first term on the right-hand side (RHS) in Eq. (1) represents the normoxic cells proliferate logistically with a rate *λ*_*n*_ when oxygen level is above the hypoxic threshold value *c*_*H*_. The second term on the RHS indicates that all hypoxic phenotypic cells switch to a normoxic phenotype when the local oxygen level exceeds *c*_*H*_. The third term indicates that normoxic cells switch to hypoxic cells when the local oxygen level is below *c*_*H*_, at a rate *µ*_*nh*_. The last term denotes normoxic cells die at a rate *γ*_*n*_ when oxygen levels drop below the necrotic threshold value *c*_*N*_.

The spatial-temporal dynamics of hypoxic cell density is given as [19]

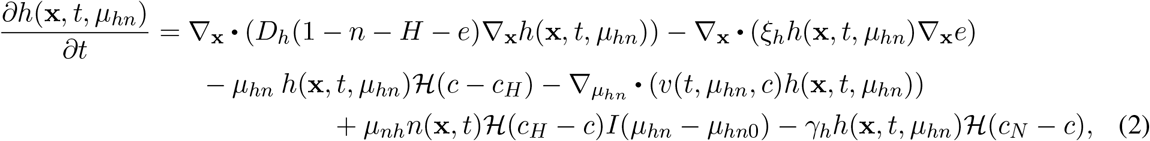

where the first term on the RHS represents the diffusion of hypoxic cells, constrained by the volumetric effects of all components in the tumor, with diffusion coefficient *D*_*h*_. The second term describes the directed movement of hypoxic cells toward regions of higher extracellular matrix (ECM) density with coefficient *ξ*, a mechanism known as haptotaxis. The third term presents hypoxic cells switch to normoxic phenotype at a rate *µ*_*hn*_ when the local oxygen level rises above *c*_*H*_. Note that this rate *µ*_*hn*_ is a dynamic quantity. Its dynamics depends on duration of hypoxic environment exposure. It is modeled by the following equation

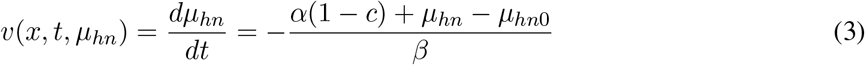

The fourth term on RHS of Eq. (2) describes hypoxic cells drift in *µ*_*hn*_ phenotype space with the velocity *v*. The velocity *v* is structured in a way that the transition rate decreases when the local oxygen concentration in the tumor decreases with respect to its maximum value 1 Here, *α* modulates the asymptotic reduction in the transition rate due to hypoxic memory. The parameter *β* regulates the rate at which *µ*_*hn*_ changes, and can be set according to the experimental observed timescales of hypoxic memory induction and withdrawal. The fifth term represents normoxic cells adopt hypoxic phenotype with a rate *µ*_*nh*_ when local oxygen levels drop below *c*_*H*_. Here, to account for the build-up of hypoxic memory, we consider that all normoxic cells first switch to the hypoxic state having the basal reverse transition rate *µ*_*hn*_ = *µ*_*hn*0_ upon future exposure to normoxia. This is formulated using the indicator function *I*(*µ*_*hn*_ *− µ*_*hn*0_), defined as follows:

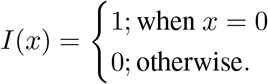

The last term on RHS in Eq. (2) defines hypoxic cells die when oxygen levels drop below *c*_*N*_.

The ECM dynamics is given as [19]

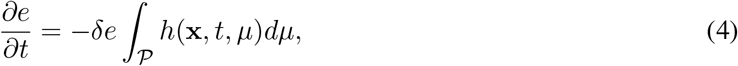

where the term on RHS represents all the hypoxic phenotype cells degrade ECM at a rate *δ*.

The spatio-temporal changes of oxygen concentration is given as

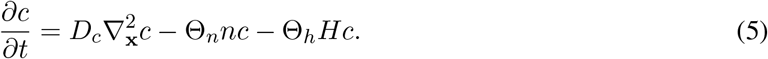

Here, the first term on the RHS of Eq. (5), oxygen is diffusing in the spatial domain with a diffusion coefficient denoted as *D*_*c*_. The second term represents the consumption of oxygen by normoxic cells at a rate of Θ_*n*_. The final term indicates that all hypoxic phenotype cells consume oxygen at the same rate, Θ_*h*_.

#### Initial and boundary conditions

For the model (Eqs. (1)-(5)), we need to prescribe the boundary conditions for Eqs. (2) and (5) as these two equations have spatial derivatives. We consider hypoxic cells not leave the one dimensional physical domain [0, *X*]. Hence, no-flux boundary condition is imposed for Eq. (2).

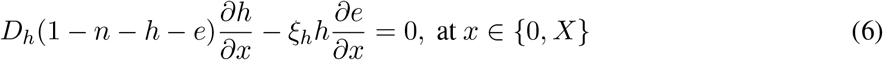

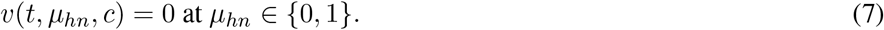

We consider oxygen is at the physical boundary is maintain a fixed level, say *c*_1_. Hence, it is given as

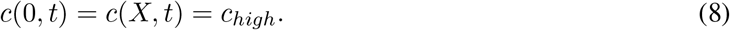

We consider a tumor located at the center of the one-dimensional physical domain [0, *X*], surrounded by the extracellular matrix (ECM). Initially, the tumor consists exclusively of normoxic cells. Furthermore, we assume that the oxygen concentration *c*_high_ is uniformly distributed throughout the entire domain. Hence the initial conditions are given as

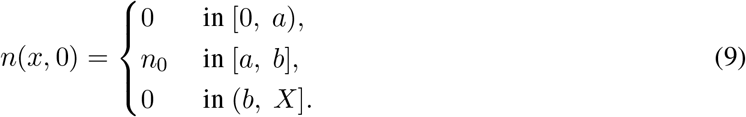

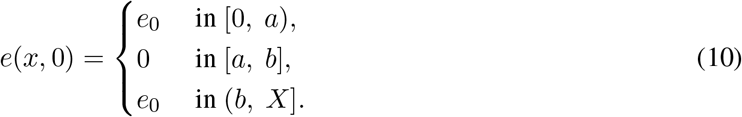

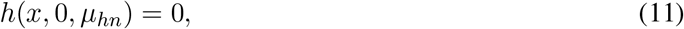

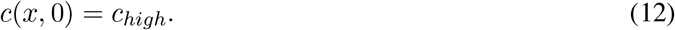

#### The protocol of oxygen levels at the tumor boundary varies over time

Here, we defined the oxygen level at the tumor boundary i.e., *x* = 0, *X* as

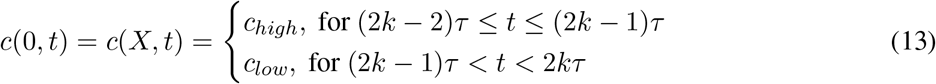

for *x* = 0, *X* and *k* = 1, 2, …, *m*. Here, 2*τ* is the period of oxygen supply at the tumor boundary and 2*τm* is the total duration of time course.

Here, we consider *n*_0_ = 0.2, *e*_0_ = 0.5, *a* = 4, *b* = 6 and *X* = 10.

#### Numerical method

To solve the proposed model, we employed the Method of Lines (MOL) in a finite difference framework. The physical domain [0, *X*] is divided into *N* − 1 intervals of uniform length 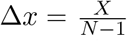,while the phenotypic domain [0, 1] is partitioned into *M* − 1 intervals of uniform length 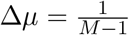 .The system of differential equations is discretized explicitly: central difference schemes are applied for spatial derivatives, while forward differences are used for temporal derivatives. The drift (convective) term in Eq. (2) is discretized using a first-order upwind scheme (Eq. (16)).

Let Δ*t* denote the time step, and let computations proceed up to final time *T*_final_, such that *p*Δ*t* = *T*_final_, where *p* is the number of time steps. We define the discrete variables as follows 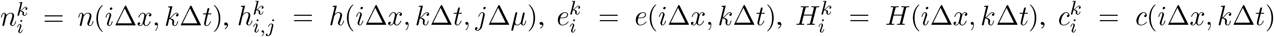and 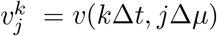 where *i* ∈ {0, 1, …, *N* −1}, *j* ∈ { 0, 1, …, *M*− 1}, and *k* ∈ {0, 1, …, *p*}.

The difference equation for Eq. (1) is obtained as,

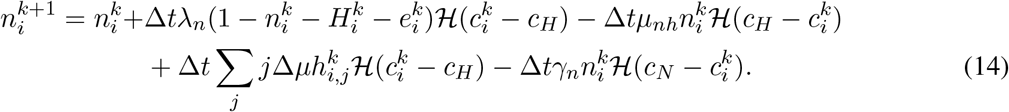

The resulted difference equation for hypoxic cells density, given by Eq. (2), is obtained as,

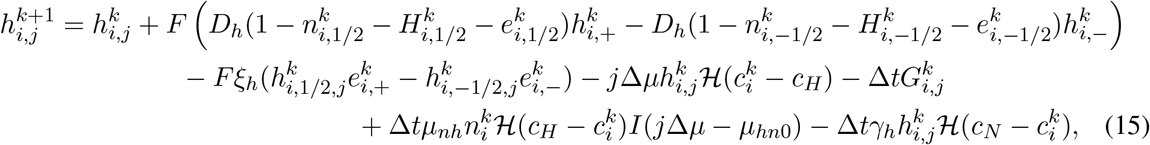

where

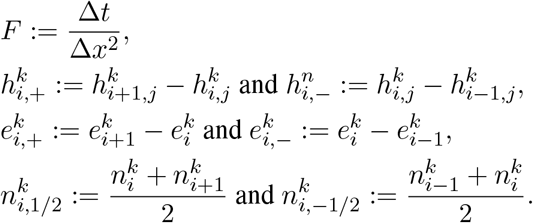

Here, 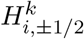 and 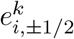 carry similar meaning.

The term 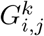 defines convection term at (*i, j*) and it is discretized using first order upwind scheme, given

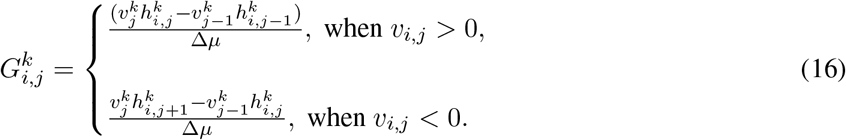

At the boundary nodes, we employ no flux boundary condition at *i* = 0 and *i* = *N* − 1 and zero convective flux at *j* = 0 and *j* = *M* −1.

The Eq. (4) is discretized as

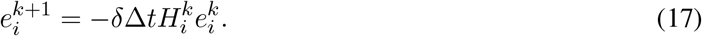

The Eq. (5) is discretized as

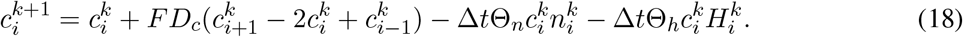

The numerical stability is ensured through the Courant–Friedrichs–Lewy (CFL) condition, which is defined as

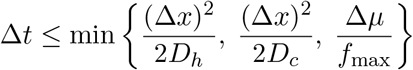

The code for this numerical exercise is available on **GitHub** link: https://github.com/gsadhuIITG/Dynamic_memory

## Author contributions

PJ, MKJ and JTG conceived and supervised research. MKJ and JTG obtained funding. GS, PJ and RKM performed research, analyzed data and wrote first draft of the manuscript. All authors contributed to reviewing and editing the manuscript.

## Supplementary Information

**Figure S1:**
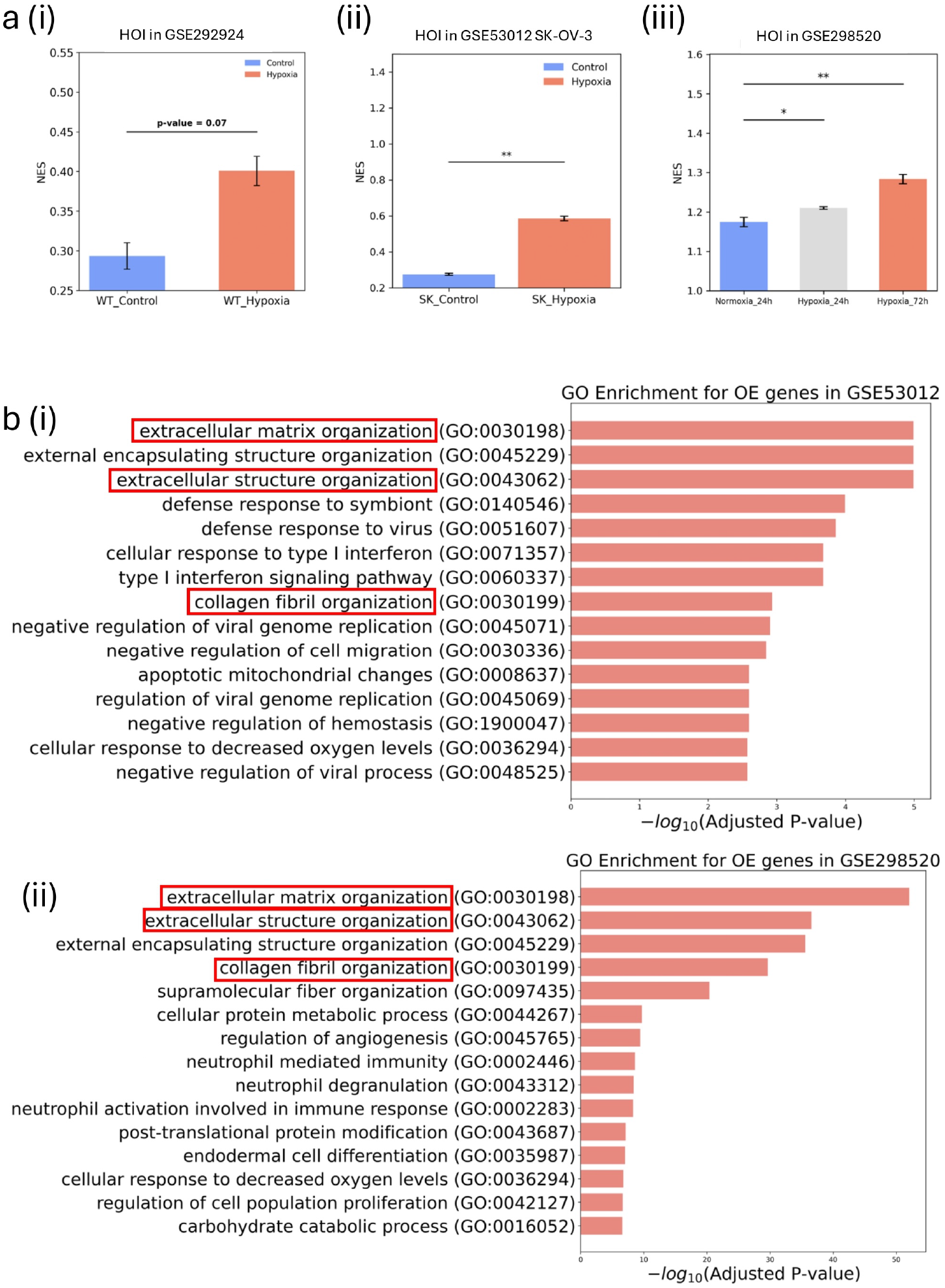
Geneset Enrichemnet Analysis and Gene Ontology analysis on publicly available cell line samples. (a) GSEA analysis for enrichment level of HOI (Hypo-methylated; Over-expressed and Invasive) geneset in (i)GSE292924, (ii) GSE53012 and (iii) GSE298520. (b) Top 15 GO classes (sorted by Adj(pvalue)) upon GO analysis of over-expressed genes in GSE53012 and GSE298520, respectively.

**Figure S2:**
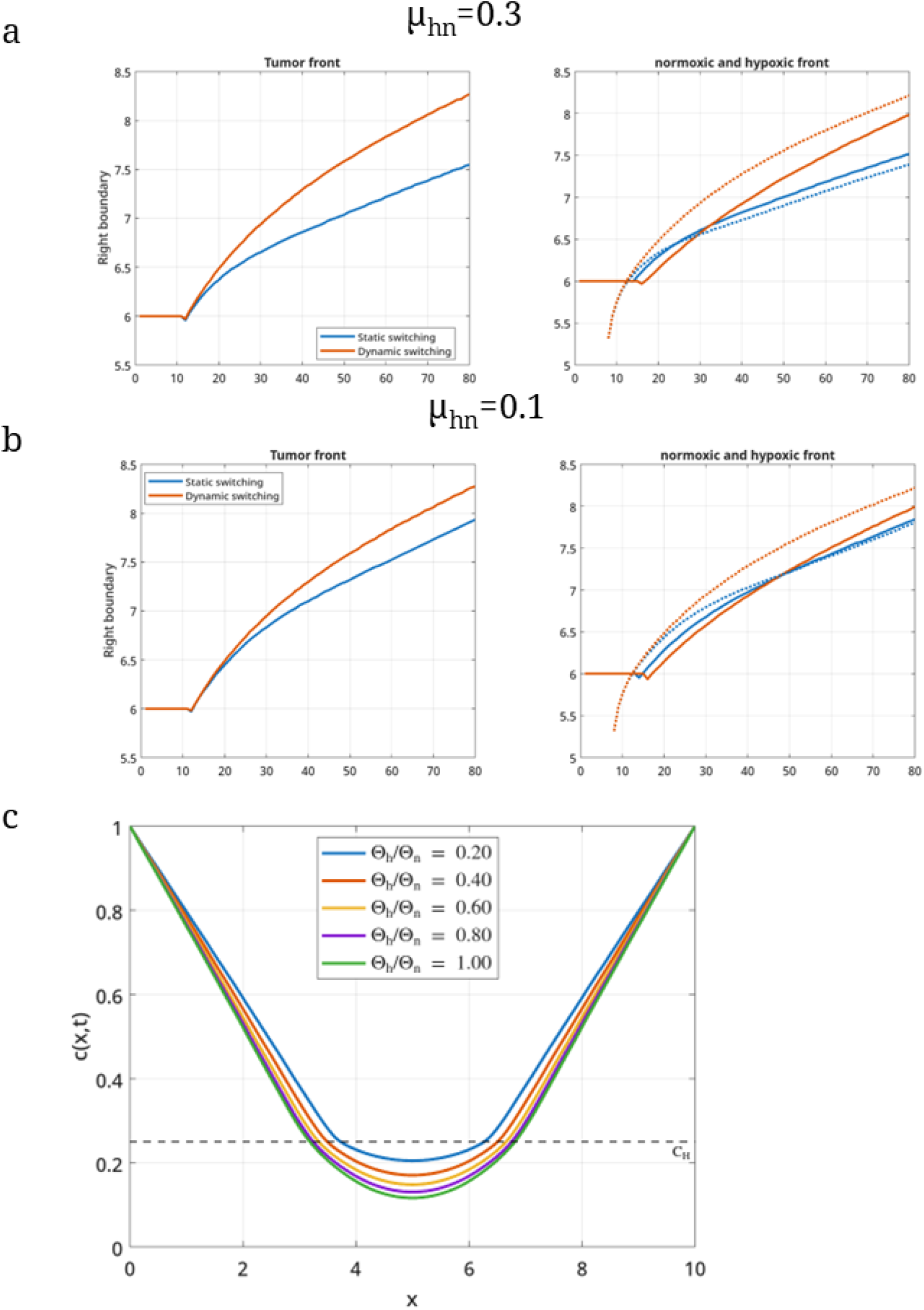
The tumor front, hypoxic and normoxic boundary propagation over time for (a) *µ*_*hn*0_ = 0.3 and (b)*µ*_*hn*0_ = 0.1 in both dynamic and fixed memory modes. (c) Oxygen profile over the spatial domain for different fold ratio of oxygen consumption rates at *t* = 60.

**Figure S3:**
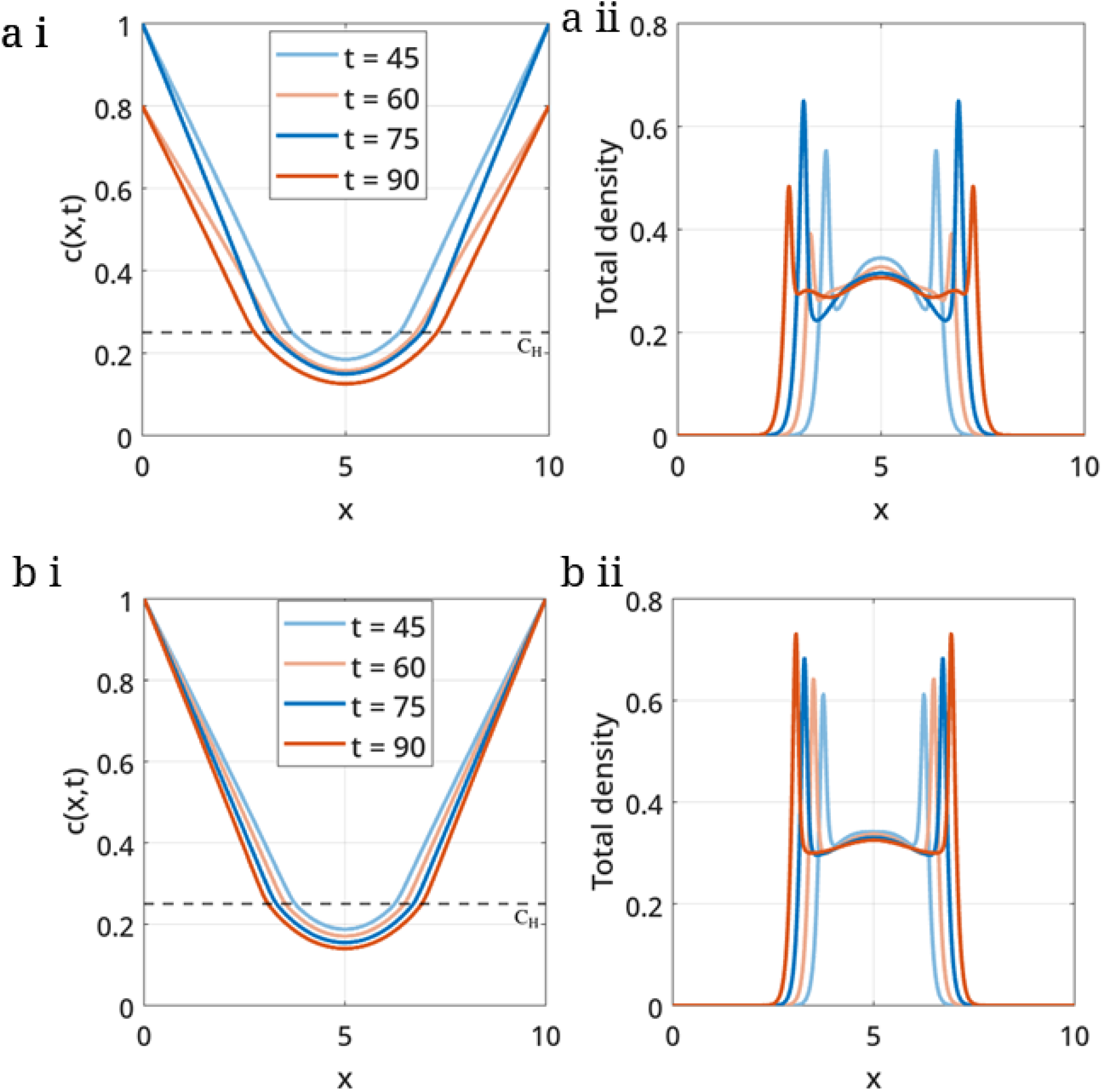
The spatial distribution of oxygen and total tumor cell density at different time points in a fixed memory regime for (a) oxygen fluctuation and (b) constant oxygen supply.

**Figure S4:**
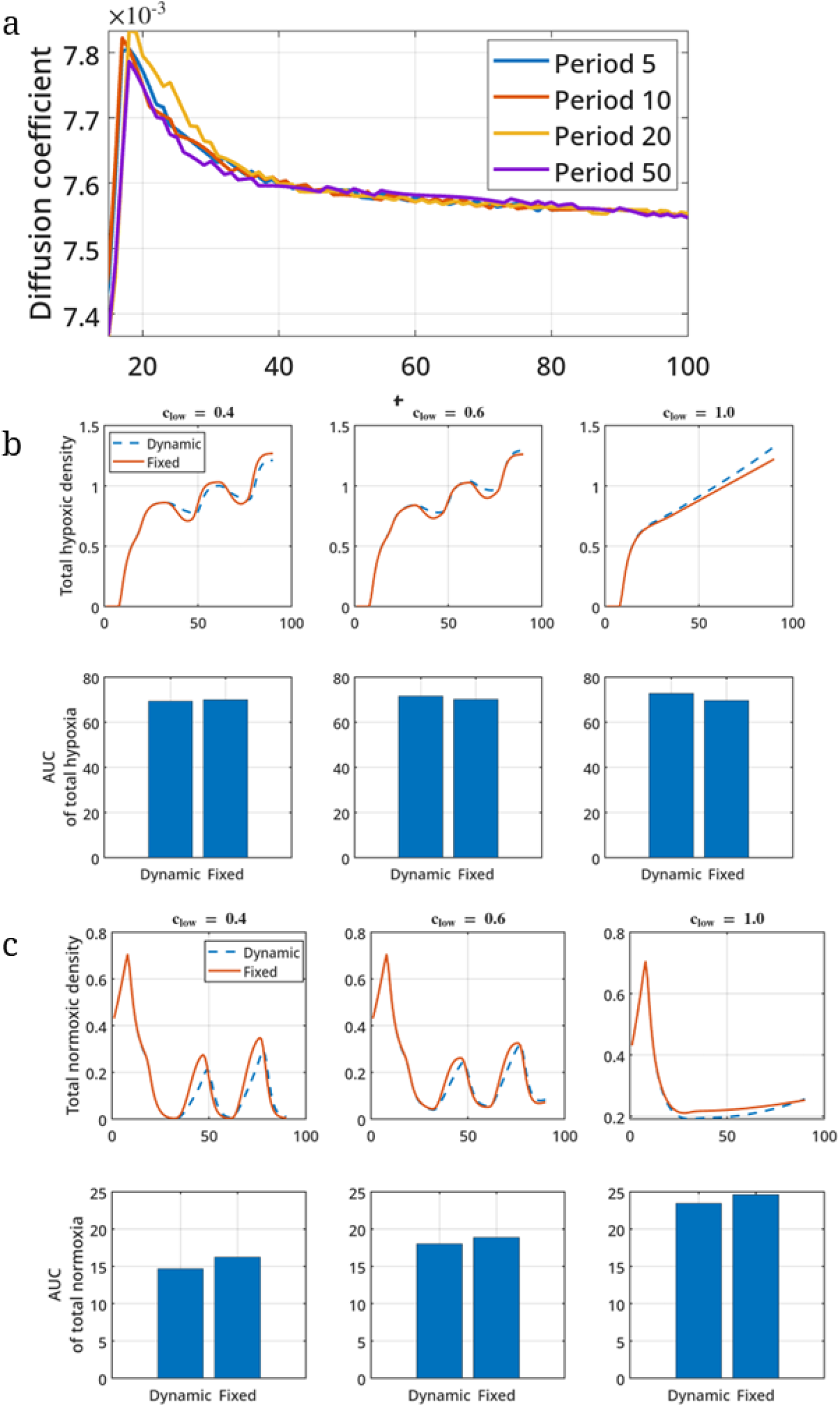
(a) Effective diffusion coefficient for different periods of oxygen fluctuation at the tumor boundary. (b) Total hypoxic cells over time and corresponding AUC for both dynamic and fixed memory modes.(c) Total normoxic cells over time and corresponding AUC for both dynamic and fixed memory modes.

**Figure S5:**
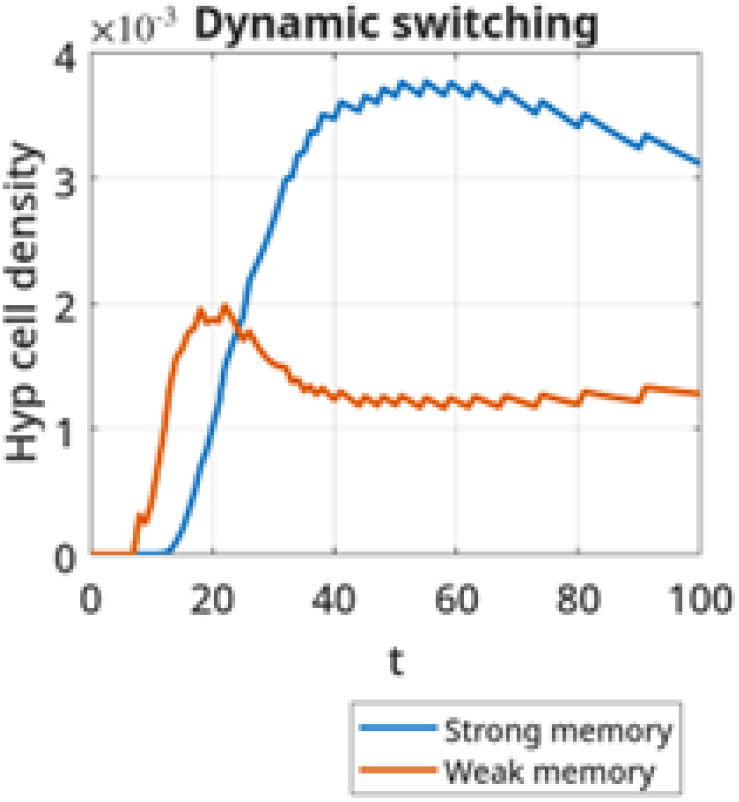
Weak memory (*µ*_*hn*_ *>* 0.25) and strong memory (0 *< µ*_*hn*_ ≤ 0.25) hypoxic cell density evolution over time in normoxic region.

## References

[1] Peter W Vaupel, Stanley Frinak, and Haim I Bicher. Heterogeneous oxygen partial pressure and ph distribution in c3h mouse mammary adenocarcinoma. Cancer Research, 41(5):2008–2013, 1981.

[2] Peter Vaupel, Michael Höckel, and Arnulf Mayer. Detection and characterization of tumor hypoxia using po2 histography. Antioxidants & redox signaling, 9(8):1221–1236, 2007.

[3] Jen-Tsan Chi, Zhen Wang, Dimitry S A Nuyten, Edwin H Rodriguez, Marci E Schaner, Ali Salim, Yun Wang, Gunnar B Kristensen, Åslaug Helland, Anne-Lise Børresen-Dale, et al. Gene expression programs in response to hypoxia: cell type specificity and prognostic significance in human cancers. PLoS medicine, 3(3):e47, 2006.

[4] Katherine L Eales, Kate ER Hollinshead, and Daniel A Tennant. Hypoxia and metabolic adaptation of cancer cells. Oncogenesis, 5(1):e190–e190, 2016.

[5] Ineŝ Godet, Harsh H Oza, Yi Shi, Natalie S Joe, Alyssa G Weinstein, Jeanette Johnson, Michael Considine, Swathi Talluri, Jingyuan Zhang, Reid Xu, et al. Hypoxia induces ros-resistant memory upon reoxygenation in vivo promoting metastasis in part via muc1-c. Nature communications, 15(1):8416, 2024.

[6] Biswanath Chatterjee, Pritha Majumder, Chun-Chang Chen, Jing-Ping Wang, Po-Hsuan Su, Hung-Cheng Lai, Ching-Chen Liu, Hsin-Nan Lin, Chen-Hsin A Yu, Hanna S Yuan, et al. Hypoxia-induced genome-wide dna demethylation by dnmt3a and emt of cancer cells. Cellular & Molecular Biology Letters, 30(1):95, 2025.

[7] Parik Kakani, Shruti Ganesh Dhamdhere, Deepak Pant, Rushikesh Joshi, Jharna Mishra, Atul Samaiya, and Sanjeev Shukla. Hypoxia-induced ctcf promotes emt in breast cancer. Cell reports, 43(7), 2024.

[8] David A Close and Paul A Johnston. Detection and impact of hypoxic regions in multicellular tumor spheroid cultures formed by head and neck squamous cell carcinoma cells lines. SLAS discovery, 27(1):39–54, 2022.

[9] José A Almeida, Diego B Avila, Gregory D Longmore, and Amit Pathak. History of hypoxia exposure aids future cell invasion according to cell type and collagen density. Molecular Biology of the Cell, 36(7):ar82, 2025.

[10] Ineŝ Godet, Yu Jung Shin, Julia A Ju, I Chae Ye, Guannan Wang, and Daniele M Gilkes. Fate-mapping post-hypoxic tumor cells reveals a ros-resistant phenotype that promotes metastasis. Nature communications, 10(1):4862, 2019.

[11] Oihana Iriondo, Desirea Mecenas, Yilin Li, Christopher R Chin, Amal Thomas, Aidan Moriarty, Rebecca Marker, Yiru J Wang, Haley Hendrick, Yonatan Amzaleg, et al. Hypoxic memory mediates prolonged tumor-intrinsic type i interferon suppression to promote breast cancer progression. Cancer research, 84(19):3141–3157, 2024.

[12] Heber L Rocha, Ineŝ Godet, Furkan Kurtoglu, John Metzcar, Kali Konstantinopoulos, Soumitra Bhoyar, Daniele M Gilkes, and Paul Macklin. A persistent invasive phenotype in post-hypoxic tumor cells is revealed by fate mapping and computational modeling. Iscience, 24(9), 2021.

[13] Roxane Verdikt and Bernard Thienpont. Epigenetic remodelling under hypoxia. In Seminars in Cancer Biology, volume 98, pages 1–10. Elsevier, 2024.

[14] Bernard Thienpont, Jessica Steinbacher, Hui Zhao, Flora D’Anna, Anna Kuchnio, Athanasios Ploumakis, Bart Ghesquière, Laurien Van Dyck, Bram Boeckx, Luc Schoonjans, et al. Tumour hypoxia causes dna hypermethylation by reducing tet activity. Nature, 537(7618):63–68, 2016.

[15] Abhishek A Chakraborty, Tuomas Laukka, Matti Myllykoski, Alison E Ringel, Matthew A Booker, Michael Y Tolstorukov, Yuzhong Jeff Meng, Samuel R Meier, Rebecca B Jennings, Amanda L Creech, et al. Histone demethylase kdm6a directly senses oxygen to control chromatin and cell fate. Science, 363(6432):1217–1222, 2019.

[16] Michael Batie, Julianty Frost, Mark Frost, James W Wilson, Pieta Schofield, and Sonia Rocha. Hypoxia induces rapid changes to histone methylation and reprograms chromatin. Science, 363(6432):1222–1226, 2019.

[17] Lacramioara Bintu. Dynamics of epigenetic regulation at the single-cell level. Biophysical Journal, 110(3):317a–318a, 2016.

[18] Adi Mukund and Lacramioara Bintu. Temporal signaling, population control, and information processing through chromatin-mediated gene regulation. Journal of theoretical biology, 535:110977, 2022.

[19] Gopinath Sadhu, Paras Jain, Jason Thomas George, and Mohit Kumar Jolly. A phenotype-structured pde framework for investigating the role of hypoxic memory on tumor invasion under cyclic hypoxia. Bulletin of Mathematical Biology, 88(2):23, 2026.

[20] Min-Zu Wu, Ya-Ping Tsai, Muh-Hwa Yang, Chi-Hung Huang, Shyue-Yih Chang, Cheng-Chi Chang, Shu-Chun Teng, and Kou-Juey Wu. Interplay between hdac3 and wdr5 is essential for hypoxia-induced epithelial-mesenchymal transition. Molecular cell, 43(5):811–822, 2011.

[21] Christina Curtis, Sohrab P Shah, Suet-Feung Chin, Gulisa Turashvili, Oscar M Rueda, Mark J Dunning, Doug Speed, Andy G Lynch, Shamith Samarajiwa, Yinyin Yuan, et al. The genomic and transcriptomic architecture of 2,000 breast tumours reveals novel subgroups. Nature, 486(7403):346–352, 2012.

[22] Aravind Subramanian, Pablo Tamayo, Vamsi K Mootha, Sayan Mukherjee, Benjamin L Ebert, Michael A Gillette, Amanda Paulovich, Scott L Pomeroy, Todd R Golub, Eric S Lander, et al. Gene set enrichment analysis: a knowledge-based approach for interpreting genome-wide expression profiles. Proceedings of the national academy of sciences, 102(43):15545–15550, 2005.

[23] Xihua Qiu, Yamin Liu, Paola Vera-Licona, Eran Agmon, Kshitiz, and Yasir Suhail. Hif-1 responsive iffls to explain specific transcriptional responses to cycling hypoxia in cancers. npj Systems Biology and Applications, 2025.

[24] Magdalena Olbryt, Anna Habryka, Sebastian Student, Michał Jarzab Tomasz Tyszkiewicz, and Katarzyna Marta Lisowska. Global gene expression profiling in three tumor cell lines subjected to experimental cycling and chronic hypoxia. PLoS One, 9(8):e105104, 2014.

[25] Song Yang, Weizhong Zhuang, Lishi Zhou, Weiwei Kong, Wanwan Zou, Qikun Zhu, Enze Bian, Bin Lin, Jianzheng Cen, Qiang Gao, et al. An engineered hypoxia-response promoter for human umbilical cord-derived mesenchymal stem cell-based therapeutics. BMC biotechnology, 25(1):59, 2025.

[26] Carine Michiels, Céline Tellier, and Olivier Feron. Cycling hypoxia: A key feature of the tumor microenvironment. Biochimica et Biophysica Acta (BBA)-Reviews on Cancer, 1866(1):76–86, 2016.

[27] Michal R Tomaszewski, Marcel Gehrung, James Joseph, Isabel Quiros-Gonzalez, Jonathan A Disselhorst, and Sarah E Bohndiek. Oxygen-enhanced and dynamic contrast-enhanced optoacoustic tomography provide surrogate biomarkers of tumor vascular function, hypoxia, and necrosis. Cancer research, 78(20):5980–5991, 2018.

[28] Siranoush Shahrzad, Kelsey Bertrand, Kanwal Minhas, and Brenda Coomber. Induction of dna hypomethylation by tumor hypoxia. Epigenetics, 2(2):119–125, 2007.

[29] Karolina Skowronski, Sonam Dubey, David I Rodenhiser, and Brenda Coomber. Ischemia dysregulates dna methyltransferases and p16ink4a methylation in human colorectal cancer cells. Epigenetics, 5(6):547–556, 2010.

[30] Evgeniy Khain, Mark Katakowski, Scott Hopkins, Alexandra Szalad, Xuguang Zheng, Feng Jiang, and Michael Chopp. Collective behavior of brain tumor cells: the role of hypoxia. Physical Review E—Statistical, Nonlinear, and Soft Matter Physics, 83(3):031920, 2011.

[31] Steffi Lehmann, Veronika Te Boekhorst, Julia Odenthal, Roberta Bianchi, Sjoerd van Helvert, Kristian Ikenberg, Olga Ilina, Szymon Stoma, Jael Xandry, Liying Jiang, et al. Hypoxia induces a hif-1-dependent transition from collective-to-amoeboid dissemination in epithelial cancer cells. Current Biology, 27(3):392–400, 2017.

[32] Kritika Saxena, Mohit Kumar Jolly, and Kuppusamy Balamurugan. Hypoxia, partial emt and collective migration: Emerging culprits in metastasis. Translational oncology, 13(11):100845, 2020.

[33] Vaishali Aggarwal, Sarthak Sahoo, Vera S Donnenberg, Priyanka Chakraborty, Mohit Kumar Jolly, and Shilpa Sant. P4ha2: A link between tumor-intrinsic hypoxia, partial emt and collective migration. Advances in cancer biology-metastasis, 5:100057, 2022.

[34] Kelsey S Johnson, Shaimaa Hussein, Priyanka Chakraborty, Arvind Muruganantham, Sheridan Mikhail, Giovanny Gonzalez, Shuxuan Song, Mohit Kumar Jolly, Michael J Toneff, Mary Lauren Benton, et al. Ctcf expression and dynamic motif accessibility modulates epithelial–mesenchymal gene expression. Cancers, 14(1):209, 2022.

[35] Alex Sigal, Ron Milo, Ariel Cohen, Naama Geva-Zatorsky, Yael Klein, Yuvalal Liron, Nitzan Rosenfeld, Tamar Danon, Natalie Perzov, and Uri Alon. Variability and memory of protein levels in human cells. Nature, 444(7119):643–646, 2006.

[36] Nathaniel A Hathaway, Oliver Bell, Courtney Hodges, Erik L Miller, Dana S Neel, and Gerald R Crabtree. Dynamics and memory of heterochromatin in living cells. Cell, 149(7):1447–1460, 2012.

[37] Litao Liu, Wenlan Liu, Lili Wang, Ting Zhu, Jianhua Zhong, and Ni Xie. Hypoxia-inducible factor 1 mediates intermittent hypoxia-induced migration of human breast cancer mda-mb-231 cells. Oncology letters, 14(6):7715–7722, 2017.

[38] Reshu Gupta, Chandramu Chetty, Praveen Bhoopathi, Sajani Lakka, Sanjeeva Mohanam, Jasti S Rao, and Dzung H Dinh. Downregulation of upa/upar inhibits intermittent hypoxia-induced epithelial-mesenchymal transition (emt) in daoy and d283 medulloblastoma cells. International journal of oncology, 38(3):733–744, 2011.

[39] Hong Liu, Feifei Jiang, Xinshan Jia, Jing Lan, Hao Guo, Erran Li, Aihui Yan, and Yan Wang. Cycling hypoxia affects cell invasion and proliferation through direct regulation of claudin1/claudin7 expression, and indirect regulation of p18 through claudin7. Oncotarget, 8(6):10298, 2016.

[40] Rob A Cairns, Tuula Kalliomaki, and Richard P Hill. Acute (cyclic) hypoxia enhances spontaneous metastasis of kht murine tumors. Cancer research, 61(24):8903–8908, 2001.

[41] Rob A Cairns and Richard P Hill. Acute hypoxia enhances spontaneous lymph node metastasis in an orthotopic murine model of human cervical carcinoma. Cancer research, 64(6):2054–2061, 2004.

[42] Einar K Rofstad, Kanthi Galappathi, Berit Mathiesen, and Else-Beate M Ruud. Fluctuating and diffusion-limited hypoxia in hypoxia-induced metastasis. Clinical cancer research, 13(7):1971–1978, 2007.

[43] Anna Chen, Jaclyn Sceneay, Nathan Gödde, Tanja Kinwel, Sunyoung Ham, Erik W Thompson, Patrick O Humbert, and Andreas Möller. Intermittent hypoxia induces a metastatic phenotype in breast cancer. Oncogene, 37(31):4214–4225, 2018.

[44] Yasir Suhail, Yamin Liu, Wenqiang Du, Junaid Afzal, Xihua Qiu, Amina Atiq, Paola Vera-Licona, Eran Agmon, and Kshitiz. Oscillatory hypoxia induced gene expression predicts low survival in human breast cancer patients. Molecular carcinogenesis, 63(12):2305–2315, 2024.

[45] Peter Vaupel and Arnulf Mayer. Hypoxia in cancer: significance and impact on clinical outcome. Cancer and Metastasis Reviews, 26(2):225–239, 2007.

[46] Fariba Tayyari, GA Nagana Gowda, Olufunmilayo F Olopade, Richard Berg, Howard H Yang, Maxwell P Lee, Wilfred F Ngwa, Suresh K Mittal, Daniel Raftery, and Sulma I Mohammed. Metabolic profiles of triple-negative and luminal a breast cancer subtypes in african-american identify key metabolic differences. Oncotarget, 9(14):11677, 2018.

[47] Lihui Liu, Jie Bai, Lanxin Hu, and Daqing Jiang. Hypoxia-mediated activation of hypoxia-inducible factor-1α in triple-negative breast cancer: A review. Medicine, 102(43):e35493, 2023.

[48] Jiapeng Yang, Yu Zhang, Shuo Wang, Peng Wang, Liang Dong, Luofei Li, Yuanqi Cheng, Xiaoyu Huang, Bin Xue, Wei Wang, et al. Dynamic rigidity changes enable rapid cell migration on soft substrates. Nature Communications, 16(1):8793, 2025.

[49] Paola Suarez-Meade, Rachel Whitehead, Steve Rosenfeld, Paula Schiapparelli, Konstantinos Konstantopoulos, and Alfredo Quinones-Hinojosa. Extracellular matrix stiffness conditions glioblastoma cells for long-term migration: Mechanical memory as a driver of invasion and recurrence in glioblastoma. Neuro-Oncology, page noaf205, 2025.

[50] EK Rofstad, K Sundfør, H Lyng, and CG Trope. Hypoxia-induced treatment failure in advanced squamous cell carcinoma of the uterine cervix is primarily due to hypoxia-induced radiation resistance rather than hypoxia-induced metastasis. British journal of cancer, 83(3):354–359, 2000.

[51] Kaitlin Graham and Evan Unger. Overcoming tumor hypoxia as a barrier to radiotherapy, chemotherapy and immunotherapy in cancer treatment. International journal of nanomedicine, pages 6049–6058, 2018.

[52] Matthew E Ritchie, Belinda Phipson, DI Wu, Yifang Hu, Charity W Law, Wei Shi, and Gordon K Smyth. limma powers differential expression analyses for rna-sequencing and microarray studies. Nucleic acids research, 43(7):e47–e47, 2015.

[53] Zhuoqing Fang, Xinyuan Liu, and Gary Peltz. Gseapy: a comprehensive package for performing gene set enrichment analysis in python. Bioinformatics, 39(1):btac757, 2023.

[54] Mark AJ Chaplain and G Lolas. Mathematical modelling of cancer invasion of tissue: dynamic heterogeneity. Networks and Heterogeneous Media, 1(3):399–439, 2006.

[55] Gopinath Sadhu and DC Dalal. Effects of non-linear interaction between oxygen and lactate on solid tumor growth under cyclic hypoxia. Bulletin of Mathematical Biology, 87(3):41, 2025.

[56] Alicia Martínez-González, Gabriel F Calvo, Luis A Pérez Romasanta, and Víctor M Pérez-García. Hypoxic cell waves around necrotic cores in glioblastoma: a biomathematical model and its therapeutic implications. Bulletin of mathematical biology, 74:2875–2896, 2012.

[57] KS Yadav and Gopinath Sadhu. Effect of inosine on recurrence of tumor after radiation therapy: A mathematical investigation. Journal of Theoretical Biology, page 112138, 2025.

